# Non-parallel transcriptional divergence during parallel adaptation

**DOI:** 10.1101/761619

**Authors:** Eva K Fischer, Youngseok Song, Kimberly A Hughes, Wen Zhou, Kim L Hoke

**Affiliations:** Department of Evolution, Ecology, and Behavior, University of Illinois, Urbana, IL 61801, USA; Department of Biology, Colorado State University, Fort Collins, CO 80523, USA; Department of Statistics, Colorado State University, Fort Collins, CO 80523, USA; Department of Biological Science, Florida State University, Tallahassee, FL 32306 USA

**Author notes:** Corresponding author: Eva K Fischer, Department of Evolution, Ecology, and Behavior, University of Illinois at Urbana-Champaign, Morrill Hall, 505 S. Goodwin Avenue, Urbana, IL 61801.

**Keywords:** *Poecilia reticulata*, transcriptomics, phenotypic plasticity, adaptation, parallel evolution

## Abstract

How underlying mechanisms bias evolution toward predictable outcomes remains an area of active debate. In this study, we leveraged phenotypic plasticity and parallel adaptation across independent lineages of Trinidadian guppies (*Poecilia reticulata*) to assess the predictability of gene expression evolution during parallel adaptation. Trinidadian guppies have repeatedly and independently adapted to high- and low-predation environments in the wild. We combined this natural experiment with a laboratory breeding design to attribute transcriptional variation to the genetic influences of population of origin and developmental plasticity in response to rearing with or without predators. We observed substantial gene expression plasticity as well as the evolution of expression plasticity itself across populations. Genes exhibiting expression plasticity *within* populations were more likely to also differ in expression *between* populations, with the direction of population differences more likely to be opposite those of plasticity. While we found more overlap than expected by chance in genes differentially expressed between high- and low-predation populations from distinct evolutionary lineages, the majority of differentially expressed genes were not shared between lineages. Our data suggest alternative transcriptional configurations associated with shared phenotypes, highlighting a role for transcriptional flexibility in the parallel phenotypic evolution of a species known for rapid adaptation.

## Introduction

Phenotypic evolution is biased by the mechanisms that link genetic variation to phenotypic variation, i.e. the genotype-phenotype map (Alberch, 1991; Pigliucci, 2010). The complex nature of the genotype-phenotype map, the existence of ‘many-to-one’ mappings across hierarchical levels of biological organization, and the multigenic nature of complex traits make it challenging to assess how mechanistic properties shape evolutionary outcomes (Boyle, Li, & Pritchard, 2017; Potticary, Morrison, & Badyaev, 2020). A deeper understanding of these phenomena will further our insight into the forces limiting and potentiating the evolution of complex traits and deepen our knowledge of how organisms respond to novel and changing environments.

Studies focusing on the independent evolution of shared traits can reveal biases in mechanisms of adaptation by asking whether similar mechanism are associated with repeated, independent evolutionary transitions. Shared phenotypes arising in response to similar selection pressures are referred to as “parallel” or “convergent” depending on the evolutionary relatedness of populations and author preference (see Glossary). Different definitions have been discussed and debated elsewhere (e.g. Arendt & Reznick, 2008; Losos, 2011; Wake, Wake, & Specht, 2011), and we use “parallel” here following the terminology conventionally applied to our study system (see below).

Understanding whether shared phenotypes also share underlying genetic and molecular mechanisms is a long-stranding focus of evolutionary biology (reviewed in Bolnick, Barrett, Oke, Rennison, & Stuart, 2018; Stern, 2013). Compelling examples of parallelism and convergence demonstrate that similar phenotypes may share underlying neural, physiological, molecular, and/or genetic mechanisms, even across highly divergent taxa (e.g. Insel and Young, 2000; Manceau et al., 2010; Pankey et al., 2014; Rosenblum et al., 2010). Similar mechanisms underlying parallel evolutionary transitions suggest the genotype-phenotype map is constrained by limits on the possible ways to construct adaptive phenotypes (Losos, 2011). In this case, phenotypic evolution may rely on a limited number of mechanistic paths, repeatedly targeting those mechanisms that yield the greatest phenotypic responses with the smallest pleiotropic costs. Indeed, the recent view of complex traits as “omnigenic” highlights the potential for near ubiquitous pleiotropy to constrain evolutionary trajectories to a very limited number of paths (Boyle et al., 2017).

In contrast, non-shared mechanisms giving rise to shared, independently derived phenotypes suggest that mechanistic flexibility may facilitate evolution by providing multiple, alternative ‘solutions’ to a given adaptive problem, thereby providing alternative paths by which organisms can reach adaptive peaks (Badyaev & Morrison, 2018; Drion, O’Leary, & Marder, 2015; Grashow, Brookings, & Marder, 2009). Empirical studies demonstrate that different mechanistic ‘solutions’ can give rise to shared phenotypes in closely related species, among populations of the same species, or even among individuals of the same population (e.g. Abouheif and Wray, 2002; Crawford and Oleksiak, 2007; Drion et al., 2015; Grashow et al., 2009; Mandic et al., 2018). Mechanistic flexibility may facilitate evolution by allowing organisms to reach the same adaptive peak from alternative starting points (e.g. distinct genetic backgrounds), and/or buffering organisms from pleiotropic effects via compensatory mechanisms that exploit the alternative paths created by many-to-one mappings. Whether the potentially “omnigenic” nature of complex traits contributes to constraints due to widespread pleiotropy or facilitates alternative paths is an area ripe for exploration (Boyle et al., 2017).

In sum, empirical evidence for both shared and distinct mechanisms underlying parallel phenotypic evolution leaves open the question of when and why either pattern dominates. Further addressing this question is a focus of theoretical and empirical work, with implications for evolutionary biology, conservation, and medicine (reviewed in Bolnick et al., 2018). In view of the potential importance of many-to-one mappings, gene expression studies provide promising starting points for surveying broadscale patterns of (non)parallelism in the mechanisms underlying complex traits. Despite this promise and substantial recent interest in (non)parallel evolution (Bolnick et al., 2018; Kaeuffer, Peichel, Bolnick, & Hendry, 2011; Oke et al., 2016; Paccard et al., 2020; Stuart et al., 2017), gene expression mechanisms underlying parallel evolution have only recently begun to be explored (Ghalambor et al., 2015; Manousaki et al., 2013; Pankey et al., 2014).

### Does phenotypic plasticity bias phenotypic evolution?

One feature of biological systems that may channel evolution into particular paths is environmentally induced phenotypic plasticity. First, both plastic and evolutionary processes may rely on a limited number of mechanistic paths, repeatedly targeting those mechanisms that yield the greatest phenotypic responses with the smallest pleiotropic costs. In this case, we expect plastic and evolutionary responses to share underlying mechanisms. Theory predicts that plastic and evolutionary changes will be in the same direction when plasticity in a novel environment is adaptive, increasing immediate survival and allowing time for evolutionary divergence via co-option of mechanisms involved in environmentally induced responses (Baldwin, 1896; Ghalambor, McKay, Carroll, & Reznick, 2007; Lande, 2009; West-Eberhard, 2003). In contrast, plastic and evolutionary changes will be in opposite directions when plasticity in a novel environment is non-adaptive, thereby increasing the strength of selection, or when plastic responses ‘overshoot’ adaptive optima and are compensated by selection (Conover, Duffy, & Hice, 2009; Ghalambor et al., 2007; Grether, 2005; Velotta & Cheviron, 2018). Importantly, plasticity may facilitate adaptation under either scenario and empirical studies document both adaptive (e.g. Fraser et al., 2014; Gleason & Burton, 2015; Li et al., 2018; Mäkinen, Papakostas, Vøllestad, Leder, & Primmer, 2016; Scoville & Pfrender, 2010; Shaw et al., 2014; Wang & Althoff, 2019) and non-adaptive (e.g. Dayan, Crawford, & Oleksiak, 2015; Ghalambor et al., 2015; W. Ho & Zhang, 2018; Pespeni et al., 2013; Schaum, Rost, Millar, & Collins, 2013) plasticity associated with adaptive evolution. Finally, plastic and evolutionary processes may rely on shared mechanistic paths because the mechanisms mediating phenotypic plasticity promote the accumulation of cryptic genetic variation that is released under new environmental conditions, thereby fostering associations between plasticity and divergence (Draghi & Whitlock, 2012; Espinosa-Soto, Martin, & Wagner, 2011).

In the present study, we combine measures of phenotypic plasticity and parallel adaptation to ask whether flexibility in underlying mechanisms shapes evolutionary trajectories. While the potentially important role of phenotypic plasticity in this process has been pointed out, few studies have explicitly considered the role of environmentally induced plasticity in parallelism (for rare examples see Oke et al., 2016; Torres-Dowdal, Handelsman, Reznick, & Ghalambor, 2012), nor asked how plasticity itself evolves (West-Eberhard, 2003). By comparing transcriptional mechanisms associated with adaptation within and across parallel lineages, we are able to assess whether plasticity biases evolutionary outcomes, to ask whether parallel phenotypic evolution is associated with shared gene expression patterns, and to speculate on the role of mechanistic flexibility in adaptive evolution.

### Guppies as a model system

Trinidadian guppies (*Poecilia reticulata*) have become a model system in ecology and evolutionary biology due to repeated, independent adaptation of natural guppy populations to distinct predation environments (Endler, 1995; Haskins, Haskins, McLaughlin, & Hewitt, 1961; D. Reznick, Butler IV, & Rodd, 2001). In Trinidad, high- and low-predation population pairs from different river drainages represent independent evolutionary lineages (Barson, Cable, & Van Oosterhout, 2009; Fraser, Künstner, Reznick, Dreyer, & Weigel, 2015; Gilliam, Fraser, & Alkins-Koo, 1993; Willing et al., 2010) in which colonization of low-predation environments has led to parallel, adaptive changes in life history traits, morphology, and behavior (Endler, 1995; Magurran, 2005; D. A. Reznick, Bryga, & Endler, 1990; D. Reznick et al., 2001; D. N. Reznick, 1997). For example, low-predation guppies shoal less tightly (Huizinga, Ghalambor, & Reznick, 2009; Magurran & Seghers, 1990b, 1991), escape more slowly (Ghalambor, Reznick, & Walker, 2004), are slower to re-commence movement following a predator encounter (Elvidge, Ramnarine, & Brown, 2014; Harris, Ramnarine, Smith, & Pettersson, 2010), and perform fewer predator inspections (Magurran & Seghers, 1990a, 1994) than their high-predation counterparts.

Building on decades of work comparing high- and low-predation populations in the wild, more recent laboratory experiments have deployed breeding designs to disentangle genetic and environmental influences of predation on phenotypic differences. These recent studies demonstrate that a combination of genetic and environmental influences shape guppy life history (Torres Dowdall et al., 2012), morphology (Fischer, Soares, Archer, Ghalambor, & Hoke, 2013; Handelsman, Ruell, Torres-Dowdall, & Ghalambor, 2014; Ruell et al., 2013; Torres-Dowdal et al., 2012), physiology (Fischer, Harris, Hofmann, & Hoke, 2014; Handelsman et al., 2013), and behavior (Fischer, Ghalambor, & Hoke, 2016b; Huizinga et al., 2009; Torres-Dowdal et al., 2012). Yet despite many years of work on whole-organism phenotypes, the mechanisms underlying adaptive phenotypic differences in guppies remain largely unexplored. A single study characterized brain gene expression in multiple populations from the same lineage during the earliest stages of adaptation (~3 generations after colonization of low-predation environments) and found a negative relationship between phenotypic plasticity and adaptive divergence (Ghalambor et al., 2015).

### Transcriptional mechanisms of plasticity, evolution, and parallel adaptation

In the present study, we take advantage of parallel phenotypic evolution in independent lineages of Trinidadian guppies (*Poecilia reticulata*) to explore patterns of flexibility and constraint in transcriptional mechanisms mediating repeated adaptation. We compare the effects of genetic background (high-versus low-predation populations) and developmental environment (rearing with and without predator cues) on brain gene expression patterns in two parallel, independent evolutionary lineages. These lineages diverged at least 600,000 years ago, with subsequent, more recent colonization of low-predation environments by high-predation ancestors within each river drainage (Fajen & Breden, 1992; Fraser et al., 2015; Willing et al., 2010). We compare whole-brain gene expression patterns *within* and *between* lineages based on evolutionary history with and rearing with predators. We focus on gene expression in the brain because the previous gene expression study in guppies used brain tissue (Ghalambor et al., 2015) and because of previous evidence for behavioral plasticity in our focal populations (Fischer et al., 2016b).

We test four hypotheses concerning the influences of underlying mechanisms on adaptive evolutionary outcomes. **Hypothesis 1** (H1), that genes with significant expression plasticity are more likely to show genetic divergence in expression; **Hypothesis 2** (H2), that the direction of plastic responses predicts the direction of genetic divergence; **Hypothesis 3** (H3), that gene expression plasticity itself evolves; and **Hypothesis 4** (H4), that parallel phenotypic adaptation across independent lineages relies on parallel gene expression changes. To test our hypotheses we (H1) quantify the extent of overlap in genes exhibiting significant expression plasticity as well as genetic expression divergence within each drainage, (H2) examine associations in the direction of developmental and genetic expression differences within each lineage, (H3) ask whether expression plasticity itself evolves, and (H4) assess whether parallel phenotypic adaptation between lineages relies on similar changes in gene expression profiles.

A number of non-mutually exclusive, interrelated predictions follow from our hypotheses. If developmental plasticity in gene expression does indeed predict genetic divergence in gene expression, we expect the magnitude (H1) and direction (H2) of gene expression plasticity to be associated with the magnitude and direction of expression divergence between high- and low-predation populations within a river drainage (i.e. within an evolutionary lineage). If gene expression plasticity itself evolves (H3), we expect the extent and/or direction of plastic responses to differ between high- and low-predation populations within a lineage. Finally, if only a few gene expression configurations (i.e. mechanistic paths) can give rise to shared adaptive phenotypes, we expect parallel adaptation to low-predation habitats to be characterized by parallel expression changes in set of genes shared between lineages (H4). In contrast, if transcriptional mechanisms are flexible, we expect gene expression divergence between high- and low-predation populations within each river to occur in largely non-overlapping gene sets. Taken together, our results allow us to assess the flexibility of transcriptional mechanisms of adaptation across timescales – from developmental plasticity, to genetic divergence within rivers, to parallel adaptation across lineages.

## Material and Methods

### Fish collection and rearing

We used a laboratory breeding design to disentangle the influences of evolutionary history with predators and developmental experience with predators on brain gene expression patterns. We established lab populations of guppies from high- and low-predation populations pairs collected from the Aripo and Quare river drainages in 2012 and 2014, respectively. To maintain the genetic variation of the original wild-caught fish, we established 20 - 30 unique family lines for each population (i.e., for each generation a single female from each family was crossed to a single male from another family) (D. N. Reznick & Bryga, 1987). We used second-generation lab born fish from these unique family lines in this study to minimize environmental and maternal effects. At birth, we split second-generation siblings into rearing environments with (pred+) or without (pred−) predator chemical cues, and they remained in these environments until the completion of the experiment (as in Fischer et al., 2016). We used only mature males in this study. To maximize the range of genetic variation captured among focal fish, all males in a given experimental group (i.e. population and rearing environment) were from distinct families; however, our split-brood design allowed us to simultaneously leverage genetic similarity among siblings and control for the effects of genetic background on developmental responses. Figure 1 provides an overview of our experimental design and interpretation of comparisons.

**Figure 1.**
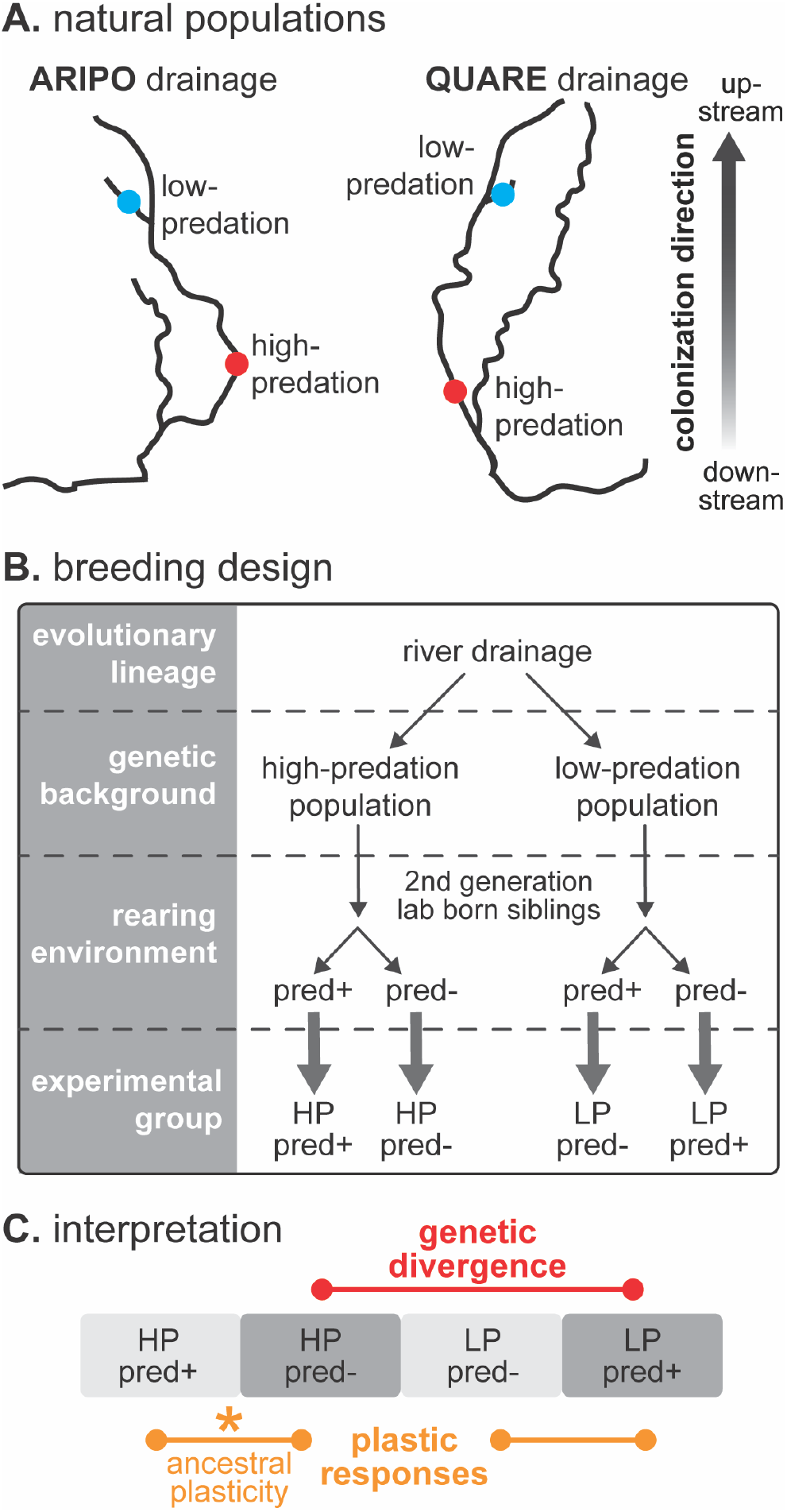
Overview of experimental design and interpretation. (A) Schematic shows Aripo and Quare river drainages in the Northern Range Mountains of Trinidad. We sampled population pairs from these two drainages, which represent distinct evolutionary lineages. (B) Wild-caught fish from each high- and low-predation population pair were reared to the second generation in the lab to control for maternal and environmental effects. At birth, second generation siblings from each population were split into rearing environments with (pred+) and without (pred−) predation cues. (C) This rearing design allows us to discern lineage effects (contrast between drainages), genetic divergence (contrast between different populations in the same environment), phenotypic plasticity (contrast between genetically similar fish in different rearing condition), and their interactions.

All guppies were individually housed in 1.5 liter tanks on a 12:12 hour light cycle at Colorado State University. Fish were fed a measured food diet once daily, receiving Tetramin™ tropical fish flake paste and hatched *Artemia* cysts on an alternating basis. Prior to tissue collection for this study, behavioral and hormone data were collected in an identical manner for all fish and these results are reported elsewhere (Fischer et al., 2016b). All experimental methods were approved by the Colorado State University Animal Care and Use Committee (Approval #12-3818A).

### Tissue collection and processing

We collected brain tissue from the Aripo and Quare lineage males described above in 2013 and 2015, respectively. To standardize any effects of recent experience, behavior, and circadian rhythm on gene expression, we collected whole brains within 10 minutes of lights-on in the morning. We collected tissue from a maximum of four fish each day, one from each experimental group per day, collected in a random order. We interpret our transcriptional data as baseline, in the sense that fish were minimally stimulated prior to tissue collection, and expression levels should therefore reflect genetic background and developmental experience more strongly than responses to immediate environmental context. Fish were anesthetized by immersion in ice water followed by rapid decapitation. Whole brains were removed, flash frozen in liquid nitrogen, and stored at −80°C until further processing. Tissue collection took <2 minutes per fish, rapid enough to minimize changes in gene expression responses to handling and dissection.

We extracted total RNA from brain tissue using the Qiagen RNeasy Lipid Tissue Mini Kit (Qiagen, Germany) following manufacturer guidelines. We prepared separate sequencing libraries for each individual using the NEBNext Ultra RNA Library Prep Kit for Illumina (New England Biolabs, Massachusetts, USA) following manufacturer instructions. Libraries were sequenced on an Illumina HiSeq 2000 at the Florida State University College of Medicine Translational Science Laboratory (Tallahassee, Florida) in May 2014 (Aripo dataset) and January 2016 (Quare dataset). For the Aripo dataset, 40 samples (N=10 per group) were combined with unique barcodes into eight samples per pool and each pool was sequenced on a single lane. For the Quare dataset, 60 samples (N=12-16 per group) were combined into three pools with 20 samples per pool and each pool was sequenced in two separate lanes. Blocks of individuals run in the same week were processed and sequenced together, and experimental groups were balanced across sequencing lanes.

### Transcript abundance estimation

To quantify gene expression, we mapped RNA sequencing reads to the guppy genome. We received 1.45 billion 100-bp paired-end reads that passed the HiSeq quality filter, averaging 14.5 million reads per sample. We used Rcorrector to amend Illumina sequencing errors (Song & Florea, 2015), and removed adapter sequences and trimmed reads for high-quality sequence using Trim Galore! (Babraham Bioinformatics, Babraham Institute). Following developer recommendations, we used a quality score of 33, a stringency of 5, and a minimum read length of 36bp. We aligned corrected, trimmed reads from both datasets to the guppy genome (http://uswest.ensembl.org/Poecilia_reticulata; downloaded November 2019) and estimated transcript abundance using STAR (Dobin et al., 2013) with default parameters. On average, 91% (range: 88.8% – 94.2%) of sequences per individual mapped to the guppy genome assembly.

### Data filtering and screening

Prior to analysis, we performed data filtering and quality screening of gene expression data. Preliminary cluster analyses revealed retinal contamination in a subset of our Aripo dataset brain samples. While opsins are expressed at low levels in the brain, the very high expression levels (>10,000 copies) in three samples pointed to retinal contamination. To deal with this issue, we devised a sample filtering and screening procedure to remove genes in which expression differences between samples were likely dominated by retinal contamination. Briefly, we first filtered genes with low expression, then we used contigs annotated as known retinal genes (Rhodopsin, red/green-sensitive opsins, blue-sensitive opsins) as seed contigs to identify other contamination-related transcripts based on high positive correlations of expression levels with seed genes. We calculated the gene-wise sum of correlations between candidate genes and seed genes and performed multiple hypothesis testing using a false discovery rate (FDR) controlling procedure. The nominal level of FDR was set to *α*=0.2 to remove presumptive contaminant contigs. Using this approach, we identified 1,559 contigs as presumptive retina-enriched genes (~ 3% of all contigs in our final assembly) which we removed from both datasets in all subsequent analyses (Table S1). More detailed descriptions of statistical procedures used for filtering are in the Supplemental Methods.

### Differential expression analysis

To test our specific hypotheses, we first needed to identify transcripts differentially expressed between experimental groups, i.e. fish with differences in evolutionary history with predators and/or developmental experience with predators (Figure 1). Due to differences in the timing of fish rearing, sample processing, and sequencing, we did not combine Aripo and Quare datasets for statistical analyses. We instead performed analysis in an identical fashion for both drainages and subsequently conducted separate analyses to test hypotheses concerning mechanistic parallelism across drainages (see below). Standard differential expression analysis packages could not accommodate the random effects in our experimental design (i.e. family and week; see below) but including these effects improved model fit and reflected our split-brood experimental design, in which siblings were split into different rearing environments at birth, and tissue was collected from fish in balanced groups of four in different weeks (see above). Fitting a generalized linear mixed model with negative binomial link to each data set, revealed that a subset of genes had statistically significant random effects due to either family, week, or both. Model comparisons between the generalized linear model without random effects and the generalized linear mixed model including week and family effects supported incorporating random effects, as likelihood ratio tests indicated better fit of the mixed models for 391 out of 19,004 genes in the Aripo dataset and 1,194 out of 19,902 genes in the Quare dataset. For consistency, we therefore performed differential expression analysis using generalized linear mixed models with random effects for all genes. Additional details of linear mixed model procedures are in the Supplemental Methods.

We normalized read counts using DESeq2 in R (Love, Huber, & Anders, 2014) and performed differential expression analysis using the lme4 package in R (github.com/lme4). Normalized count data were modeled using a generalized linear mixed model with negative binomial distribution as described above. We included population of origin, rearing environment, and their interaction as fixed effects. In addition, we included family (siblings split across rearing environments) and week (tissue was collected from animals in balanced blocks across multiple weeks) as random effects. A Wald’s test provided p-values for main effects and interaction effects for each gene (Lehmann & Romano, 2005). We adjusted p-values for multiple hypothesis testing using a direct approach for FDR correction (Storey, 2002) as implemented in the fdrtool package in R (Strimmer, 2008). We considered transcripts differentially expressed (DE) if the adjusted p-value for an effect was below 0.05.

To examine whether differential expression calls were influenced by transcript abundance, we compared mean and median counts of DE and non-DE genes using two sample t-tests and Wilcoxon rank sum tests. We also compared overall gene-wise variance between DE and non-DE groups with respect to main effects of population of origin or rearing environment using Siegel-Tukey tests and Kolmogorov-Smirnov tests. As sample sizes differed between our two datasets, we also performed a down-sampling simulation to test for the effects of sample size on DE gene number (Supplemental Methods).

We performed GO term enrichment analysis for all sets of DE transcripts using ‘Biological Processes’ in the topGO package in R (Alexa & Rahnenfuhrer, 2016) and GO annotation information from the guppy genome (http://uswest.ensembl.org/Poecilia_reticulata; downloaded November 2019).

### Statistical tests of specific hypotheses

To test Hypothesis 1, we evaluated overlap in transcripts differentially expressed between populations (i.e. population DE genes) and between rearing conditions (i.e. rearing DE genes). To test the hypothesis that initial plastic responses following colonization of low-predation environments might drive subsequent divergence, we specifically compared genes with significant ancestral plasticity (i.e. differences between high-predation siblings reared with and without predators; see Glossary and Figure 1) to genes with significant genetic expression divergence (high- versus low-predation fish reared in the derived, predator-free environment) within each drainage. We used FDR-corrected post-hoc tests of simple effects to identify DE genes in these comparisons and estimate log-fold gene expression changes.

To test Hypothesis 2, we compared the association between directions of plasticity and divergence (concordant vs. non-concordant) in (1) those genes with *both* significant ancestral plasticity and significant population differences, and (2) *all* transcripts with significant population differences in expression, including those without significant expression plasticity. The second comparison addressed the possibility that even subthreshold expression plasticity in response to rearing environment could influence patterns of expression divergence, especially given the high false negative rate in our conservative DE analyses. We used log-fold expression changes to determine whether gene expression differences were in the same/concordant direction (i.e. upregulated in high-predation populations and in response to rearing with predator cues, or down-regulated in high-predation populations and in response to rearing with predator cues) or opposite/non-concordant direction (upregulated in high-predation populations but down-regulated in response to rearing with predators, or vice versa).

Statistical approaches to test Hypotheses 1 and 2 that compare gene-wise contrasts with respect to a common experimental group (high predation fish reared without exposure to predators) are biased and nontrivial to correct, (for illustration of this point see a classical example in the Supplementary Methods and the discussion by Mallard, Jakšić, & Schlötterer, 2018). We therefore instead evaluated whether significant effects of rearing condition in the ancestral population tended to be associated with significant population effects across all genes rather than evaluating individual genes. To do so, we focused on (a) the probability of significant population simple effects conditional on significant rearing effects in the high-predation population; and (b) the probability of significant population simple effects conditional on non-significant rearing effects (Cao, Zhou, Breidt, & Peers, 2020; Hwang & Liu, 2010; Storey, 2007). We tested whether the conditional probabilities of gene expression divergence between high- and low-predation populations differed for genes that exhibited ancestral plasticity and those that did not (Hypothesis 1). We further tested whether conditional probabilities for upregulated or downregulated genes in the low-predation population compared to the high-predation population differed depending on the direction of plasticity (Hypothesis 2). To draw inference on these conditional probabilities across all genes, we adapted a parametric bootstrap method (Efron & Tibshirani, 1993) similar to the approach implemented by (W. C. Ho & Zhang, 2019). As the number of genes is large, the estimated conditional probabilities are accurate and have statistical properties that make the use of parametric bootstrap valid. We calculated parametric bootstrap confidence intervals for these differences in conditional probabilities as detailed in the Supplemental Methods, with confidence intervals that do not include zero indicating a significant association between gene expression divergence and plasticity.

To test Hypothesis 3, we examined the subset of transcripts with significant interaction effects in our differential expression analysis (see above). We grouped genes with significant interaction effects into categories for the evolution of plasticity outlined by Renn and Schumer (2013) based on *post hoc* differences between rearing environments within a population (p<0.05): (1) Assimilated: plasticity in the ancestral high-predation population but a loss of plasticity in the derived low-predation population; (2) Accommodated: a change in the degree, but not the direction, of plasticity in the derived as compared to the ancestral population; (3) Reversed: opposing directions of plasticity in high-versus low-predation populations; (4) Evolved plastic: no plasticity in the ancestral high-predation population but an emergence of plasticity in the derived low-predation population; and (5) Unclassified: genes that had a significant interaction effect but no significant *post hoc* rearing differences and therefore could not be unambiguously classified into one of the above categories. All statistical tests and data visualization were performed in R (version 3.5.1; The R Foundation for Statistical Computing).

To test Hypothesis 4, we evaluated overlap in DE transcript identity and expression direction between the Aripo and Quare lineages. We used chi-square tests of independence to test for greater than chance overlap *between* drainages in those gene sets differentially expressed based on rearing environment or population of origin *within* drainages. We also used chi-square tests to test for biases in concordant versus non-concordant expression between drainages. We performed two additional comparisons to address potential bias in our conclusions due to high false negative rates in gene expression data (Rice, Schork, & Rao, 2008). First, we performed the same analysis as before at two less stringent p-value cut-offs (p=0.1 and 0.2). Second, for comparisons of expression direction, we made an additional, more conservative comparison by including not only those genes differentially expressed in *both* drainages (i.e. the intersection of DE gene lists), but also those genes differentially expressed in *either* drainage (i.e. the union of DE gene lists) in the analysis. We again used log-fold expression differences to call gene expression as being in the same/concordant or opposite/non-concordant direction between drainages.

## Results

### Differential gene expression and GO term enrichment

In order to test our specific hypotheses, we first summarized differential expression patterns in both datasets. In the Aripo drainage, 766 genes were differentially expressed (DE) between high- and low-predation populations, 1,118 genes were differentially expressed between pred- and pred+ fish, and 825 genes had interaction effects (Fig. 2; Table S2). DE genes were enriched for GO categories including DNA integration, immune function, and morphogenesis. In the Quare drainage, 3,831 genes were differentially expressed between high- and low-predation populations, 791 genes were differentially expressed between pred- and pred+ fish, and 586 genes had interaction effects (Fig. 2; Table S3). DE genes were enriched for GO categories including immune function, developmental processes, and neurogenesis. Complete results of GO enrichment analyses for population of origin, rearing, and interaction effects are in the Supplemental Information (Tables S4 & S5).

**Figure 2.**
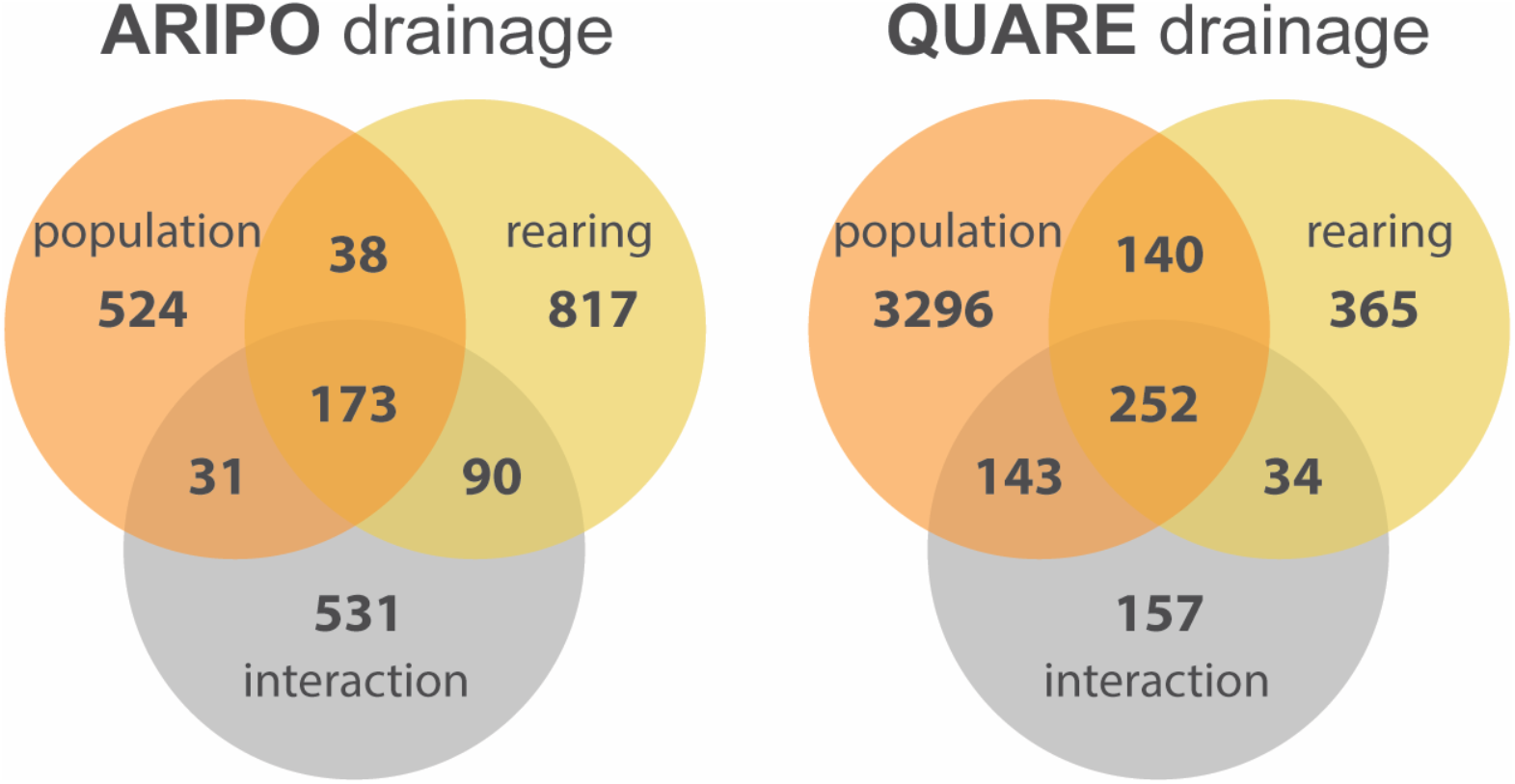
Summary of differential expression analyses. We identified many transcripts in both the Aripo (left) and Quare (right) drainages whose expression levels differed based on evolutionary history of predation (population of origin; orange), developmental experience of predation (rearing; yellow), and their interaction (grey).

Given the marked difference in the total number of DE genes between datasets, we looked for evidence that this pattern might arise from technical variation, expression bias, or differences in sample size between the two datasets. While we cannot rule out some influence of technical variation, we found no evidence for differences in sequence quality, read alignment statistics, or variance in transcript abundance to suggest that variation in the number of DE transcripts between drainages was of technical origin. Nor did we find evidence that identification of DE transcripts was biased by transcript expression level: mean and median gene expression were significantly lower in DE (population, rearing, and/or interaction effects) as compared to non-DE transcripts in both the Aripo (Fig. S1; mean: DE=263, non-DE=344, p<0.0001; median: DE=65, non-DE=107, p<0.0001), and Quare (Fig. S1; mean: DE=264, non-DE=334, p<0.0001; median: DE=86, non-DE=101, p<0.0001) datasets. A down-sampling simulation provided no evidence for a reliance of the greater number of population DE genes in the Quare dataset on the larger sample size of this dataset (Figure S2). In sum, these tests suggest that identification of DE transcripts was not influenced by relatively greater expression magnitude or variance of these transcripts, nor by sample size differences.

### Expression plasticity biases the propensity for genetic expression divergence

Evolutionary theory and empirical studies suggest that environmentally induced plasticity may facilitate adaptation to novel environments. To test Hypothesis 1, we asked whether genes with significant ancestral expression plasticity were more likely to show genetic divergence in expression. We examined the extent of overlap in gene sets (1) differentially expressed between high-predation fish reared with and without predators, to those (2) differentially expressed between populations adapted to high-versus low-predation environments. In both drainages, the DE genes with significant ancestral plasticity were more likely to show significant population divergence than expected by chance (Aripo: 264 transcripts, 95% CI for differences in conditional probability of significant divergence for plastic genes compared to conditional probability of divergence for non-plastic genes: [0.125, 0.150]; Quare: 409 transcripts, 95% CI: [0.211,0.239]), supporting an association between plastic responses and genetic divergence.

### Expression plasticity biases the direction of genetic expression divergence

To test Hypothesis 2, we asked whether the directions of expression plasticity and genetic expression divergence were associated. We did so for those genes with both significant ancestral plasticity and population differences, as well as the larger set of all transcripts with significant population differences (i.e. including those with and without significant rearing differences). We performed the second, more conservative comparison because we reasoned that even subthreshold expression plasticity could influence the direction of adaptive divergence. Within both drainages, we found a signature of non-adaptive plasticity associated with genetic expression divergence. Approximately 84% of DE genes exhibited population and rearing expression changes in non-concordant directions in the Aripo drainage (95% CI for difference in probability of same versus opposite expression direction: [−0.564,−0.499]), and 68% of DE genes exhibiting non-concordant expression in the Quare drainage (95% CI: [−0.520,−0.456]) (Fig. 3). Patterns were consistent in both drainages when we considered the larger set of all genes with significant population expression differences (Aripo: 80% of genes with non-concordant expression, 95% CI: [−0.811,−0.752]; Quare: 66% of genes with non-concordant expression, 95% CI: [−0.803,−0.733]).

**Figure 3.**
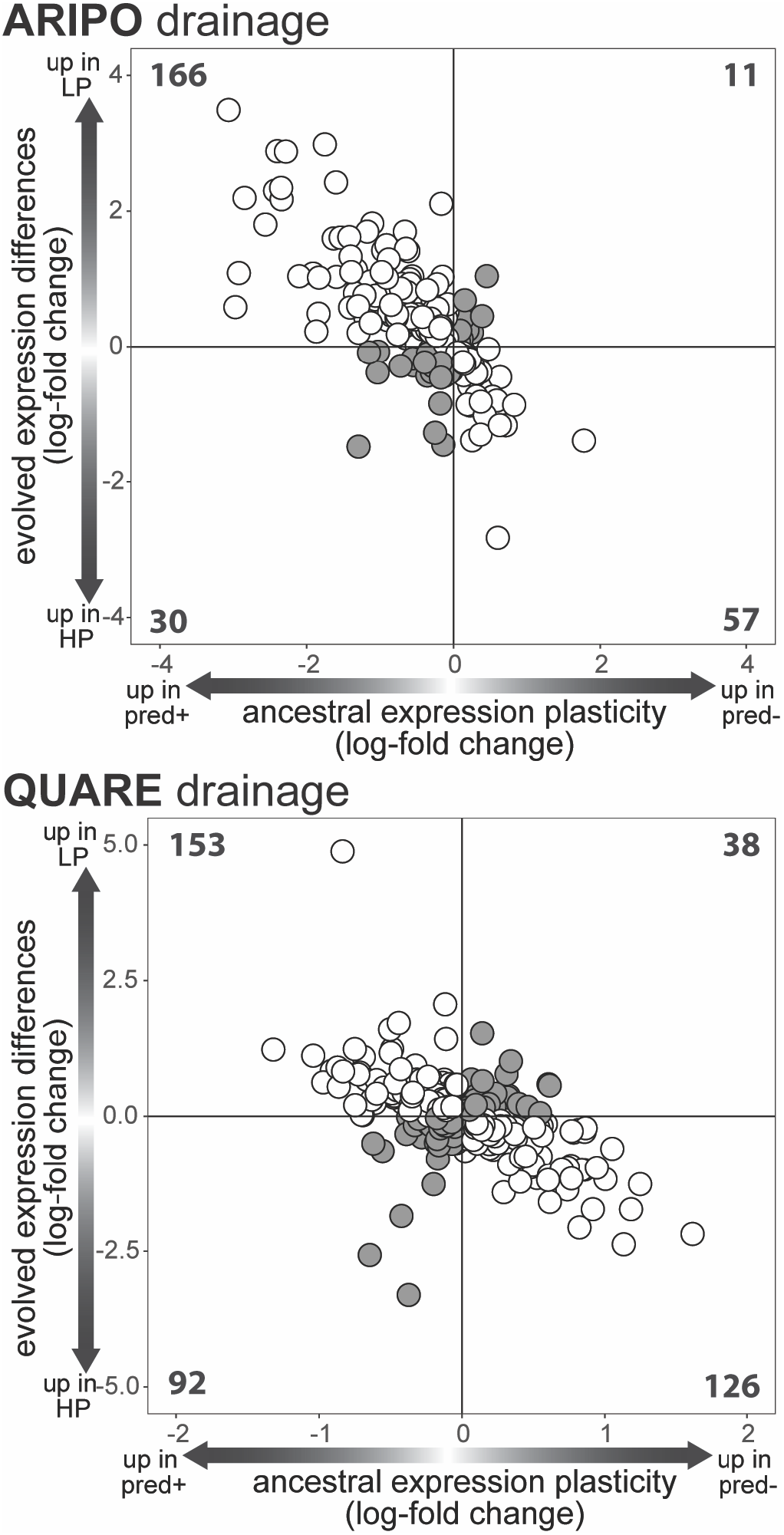
Relationship between ancestral plasticity and evolved expression divergence. Genes exhibiting significant ancestral expression plasticity (i.e. different between high-predation fish reared with and without predator cues) were more likely to also differ in expression between high- and low-predation populations reared without predator cue exposure in both the Aripo (top) and Quare (bottom) lineages. Each circle represents expression differences in a single transcript with statistically significant ancestral plasticity in the high-predation population as well as significant expression differences between high- and low-predation populations. Plastic expression changes were more likely to be in the opposite (white circles) than in the same (grey circles) direction as genetic expression divergence. The number of transcripts in each quadrant is indicated on the graphs. Note that axes differ between plots in order to best visualize variation within each dataset.

### Evolution of expression plasticity

We next explored the evolution of expression plasticity itself to test Hypothesis 3. We identified genes with evolved expression plasticity as those in which the direction and/or degree of rearing effects depended on population of origin, i.e. those genes with significant population-by-rearing interaction effects. The number of genes with significant expression plasticity that evolved (i.e. significant rearing and interaction effects) was approximately a quarter to a third of the number of genes with significant expression plasticity that did not evolve (i.e. significant rearing effect but no interaction; 263 (24%) in Aripo and 286 (36%) in Quare) (Fig. 2).

To further characterize how expression plasticity evolved, we subdivided genes with significant population-by-rearing interaction effects into one of five categories: assimilated, accommodated, reversed, evolved plastic, or unclassified (Fig. 4A; based on Renn and Schumer, 2013). We found many transcripts that exhibited evolution of plasticity, with all five categories represented in both datasets (Fig. 4B). Of the genes with significant interaction effects, in the Aripo drainage 263 (32%) transcripts showed patterns of expression assimilation, 81 (10%) transcripts showed patterns of expression accommodation, 113 (14%) transcripts exhibited reversed plasticity, and 182 (22%) transcripts evolved plasticity in the derived population. In the Quare drainage, 164 (28%) transcripts showed patterns of expression assimilation, 110 (19%) transcripts showed patterns of expression accommodation, 128 (22%) transcripts exhibited reversed plasticity, and 95 (16%) transcripts evolved plasticity in the derived population (Fig. 4B).

**Figure 4.**
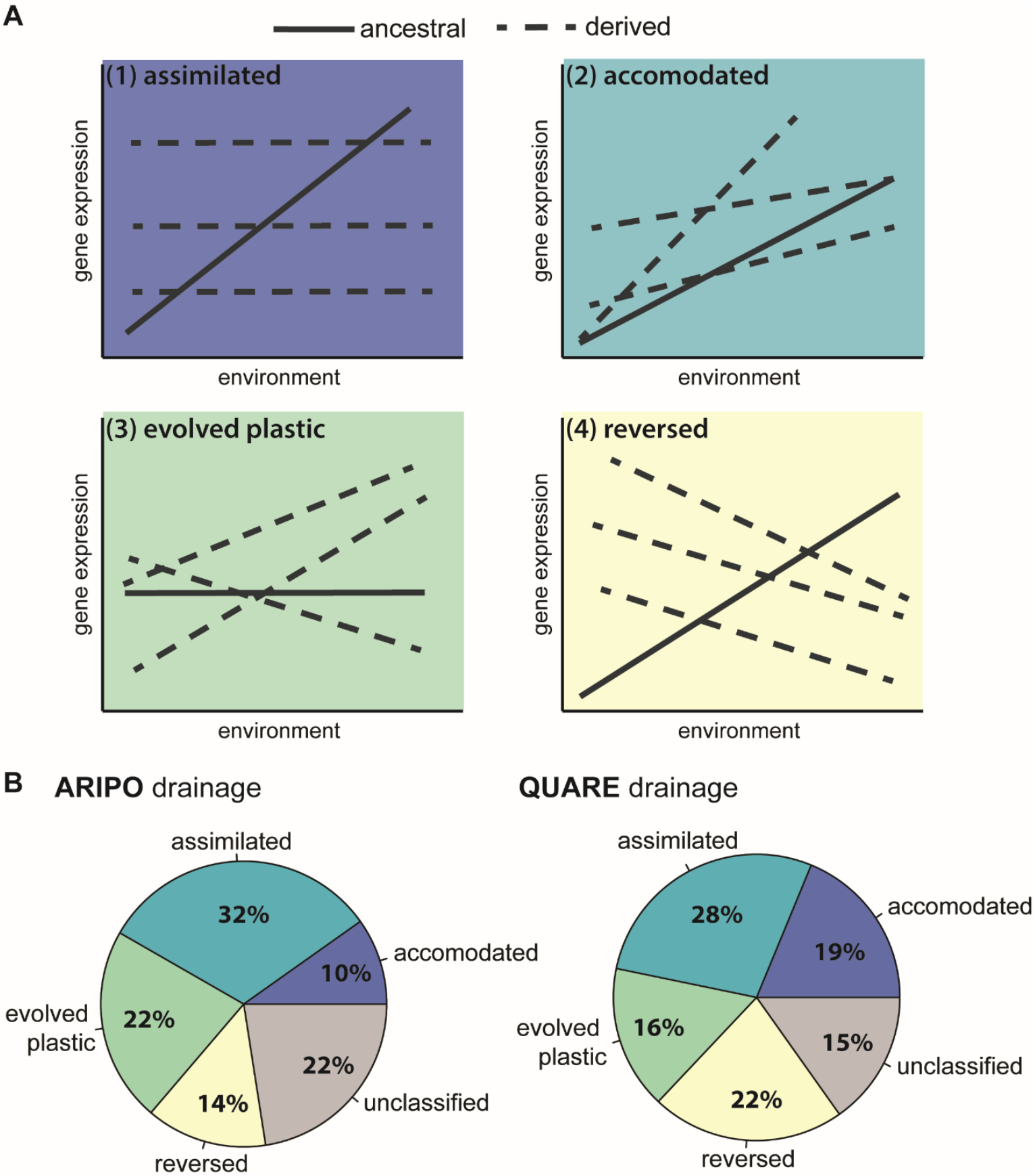
Evolution of transcript expression plasticity. (A) We grouped transcripts with significant interaction effects into one of four categories based on patterns of evolution in expression plasticity: (1) Assimilated: plasticity in the ancestral high-predation population but a loss of plasticity in the derived low-predation population; (2) Accommodated: a change in the degree, but no the direction, of plasticity in the derived as compared to the ancestral population; (3) Evolved plastic: no plasticity in the ancestral high-predation population but an emergence of plasticity in the derived low-predation population; (4) Reversed: opposing directions of plasticity in high-versus low-predation populations. We categorized remaining transcripts that had a significant main interaction effect, but no significant *post hoc* rearing differences as unclassified. Adapted from Renn & Schumer (2013). (B) All categories were represented in both Aripo and Quare datasets.

### Parallelism in gene expression patterns across drainages

To test Hypothesis 4 that shared mechanisms mediate parallel phenotypic adaptation across drainages, we asked whether differentially expressed genes *within* drainages were overlapping in identity and expression direction *between* drainages. Of the genes that diverged between high- and low-predation populations within a drainage (i.e., population main effect), 174 were overlapping between drainages (Table S6), more than expected by chance (X^2^=8.02, p=0.0046). However, the direction of expression divergence among these overlapping genes was not associated across the two drainages (X^2^=0.16, p=0.686): 91 genes (52%) had concordant expression changes and 83 genes (48%) had non-concordant expression changes between drainages (Fig. 5). To control for the influence of false negatives on our conclusions, we repeated this comparison at less stringent significance adjusted p-value cut-offs. At a p<0.1, we found marginally significant overlap in gene identify (347 overlapping genes; X^2^=3.38, p=0.06), but no association in expression direction (X^2^=0.65, p=0.42), and at p<0.2 we found no significant overlap or association in expression direction. These patterns remained unchanged when we restricted the analysis to those genes with strongest evidence for adaptive divergence (see Methods and Table S8).

**Figure 5.**
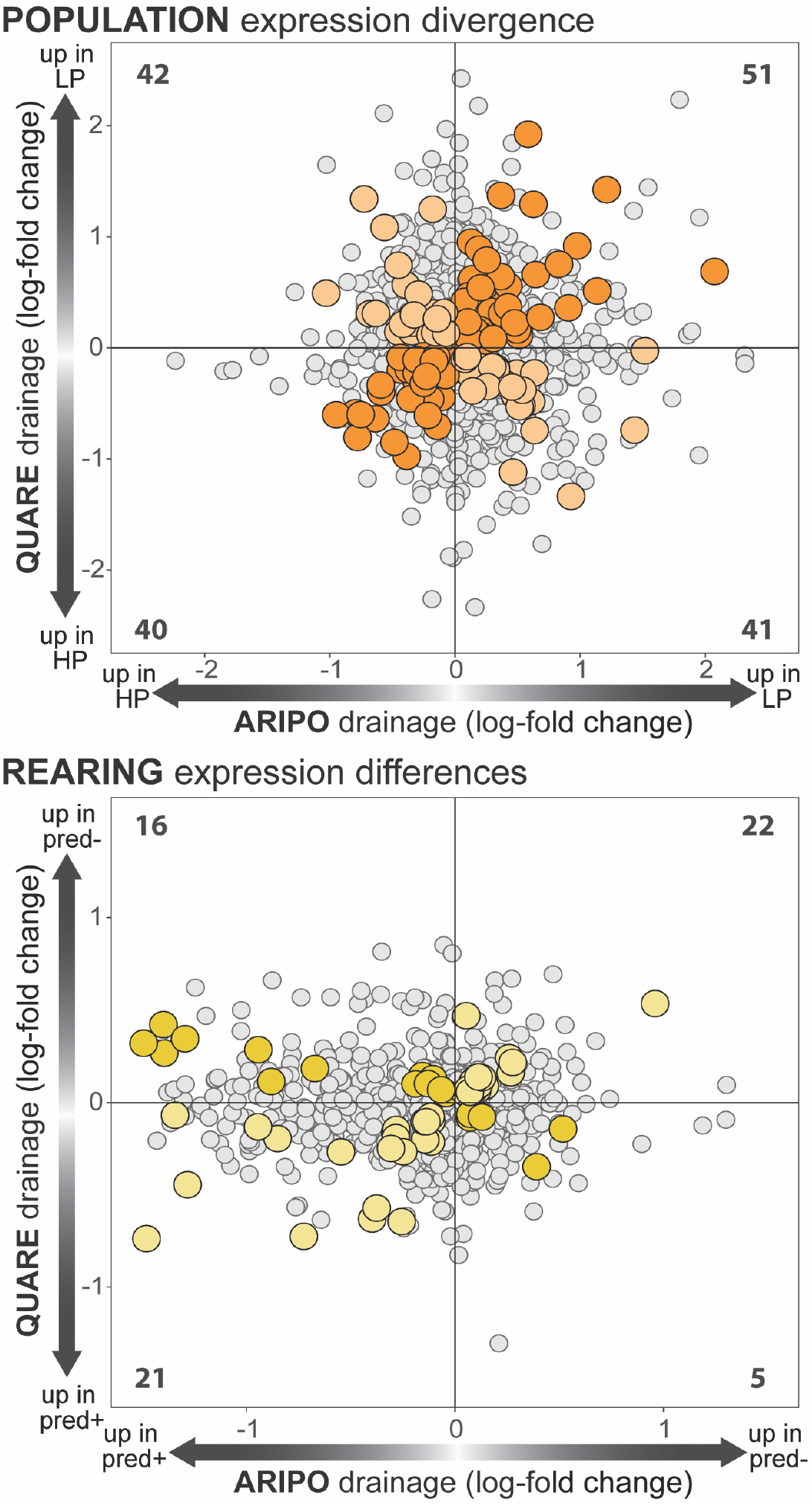
Overlap in population and rearing expression differences across evolutionary lineages. Of the transcripts differentially expressed based on population of origin (top), 124 were overlapping between drainages. Of these, 66 were concordantly differentially expressed (i.e. expression divergence in the same direction in both drainages; dark orange circles) and 58 were not (i.e. expression divergence in opposite directions between drainages; light orange circles), no different than expected by chance. Of the transcripts differentially expressed based on rearing environment (bottom), 35 were overlapping in the two drainages. Of these, 19 were concordantly differentially expressed (dark yellow circles) and six were not (light yellow circles), more non-concordance than expected by chance. The number of differentially expressed transcripts in each quadrant is indicated on the graphs and transcripts differentially expressed in one but not both drainages are shown in grey.

Of the genes that showed significant expression plasticity (i.e. main effect of rearing), 64 transcripts were overlapping between drainages (Table S7), more than expected by chance (X^2^=14.06, p=0.0001). Of these 43 (67%) had plastic changes in the same direction (i.e. concordant) and 21 (33%) had expression changes in opposite directions (i.e. non-concordant) between drainages, a stronger signature of concordant expression between river drainages than expected by chance (X^2^=7.56, p=0.006) (Fig. 5). At less stringent p-value cut-offs for differential expression we found no more overlap in gene identity than expected by chance (p<0.1: 108 overlapping genes; χ^2^=0.05, p=0.81; p<0.2: 276 overlapping genes; X^2^=1.36, p=0.24).

## Discussion

We examined transcriptomic differences associated with adaptive divergence and rearing environment across repeated, independent evolutionary lineages in guppies. Within lineages, we observed phenotypic plasticity in gene expression patterns as well as the evolution of gene expression plasticity itself. Plastic genes were more likely to exhibit population differences in expression, and these population differences were more likely to be in the opposite, rather than the same, direction as ancestral expression plasticity. Comparing across lineages, DE gene sets for population and rearing within each lineage were largely non-overlapping (>75% non-shared identity) between lineages. Nonetheless, the small number of DE genes that did overlap between drainages was more than expected by chance. Taken together, our findings suggest that a small number of core genes may be repeatedly targeted during colonization of low-predation environments, but that largely non-shared, alternative transcriptional solutions are associated with parallel phenotypic adaptation across lineages. Combining studies of phenotypic plasticity and parallel adaptation, we highlight a role for transcriptional flexibility in the parallel phenotypic evolution of a species known for rapid adaptation.

### Impacts of gene expression plasticity on expression evolution

A growing number of studies are using transcriptomic and proteomic surveys to address the long-standing debate on the role of plasticity in evolution, with contrasting results favoring the alternative hypotheses that adaptive (Fraser et al., 2014; Gleason & Burton, 2015; Li et al., 2018; Mäkinen et al., 2016; Scoville & Pfrender, 2010; Shaw et al., 2014; Wang & Althoff, 2019) versus non-adaptive (Dayan et al., 2015; Ghalambor et al., 2015; W. Ho & Zhang, 2018; Pespeni et al., 2013; Schaum et al., 2013) plasticity facilitates adaptation. We recently proposed that non-adaptive plasticity dominates during the earliest stages of rapid evolution, and that adaptive plasticity may contribute to subsequent fine-tuning of phenotypes (Fischer et al., 2016a). In line with this prediction and our previous findings, we report here that the strong signature of non-adaptive plasticity in brain gene expression (89% of transcripts) observed in guppy populations in the earliest stages of adaptation to low-predation environments (Ghalambor et al., 2015) is present but weaker in long-established low-predation populations. Furthermore, in the current study, this signature of non-adaptive plasticity was more apparent in the more recently diverged Aripo lineage (80% of genes) as compared to the older Quare lineage (65% of genes). Our findings highlight the need to explicitly consider how plasticity relates to divergence throughout successive stages of adaptation in order to build a more holistic understanding of the role of plasticity in adaptation.

### Evolution of plasticity in gene expression

Diverse transcriptional patterns could accompany evolved differences in behavioral plasticity (Renn & Schumer, 2013), but relevant data characterizing the evolution of gene expression plasticity has been lacking. We found that plasticity itself evolved in approximately one third of genes with significant ancestral expression plasticity in both lineages. As was observed for transcriptomic evolution of gill tissue in stickleback fish (Gibbons, Metzger, Healy, & Schulte, 2017), plasticity evolution showed no consistent pattern, with genes gaining, losing, and switching the direction of expression plasticity between ancestral and derived populations in both lineages. The lack of consistency in these patterns and the dearth of studies that have characterized the evolution of gene expression plasticity make these patterns difficult to interpret at present. Some of the diversity is likely associated with adaptive phenotypic divergence between high- and low-predation populations, in which initially plastic behavioral shifts may become fixed, eliminated, or altered over time. For example, evolution in gene expression plasticity could reflect the gains, losses, and switches behavioral plasticity in these populations (Fischer et al., 2016b). Alternatively, compensatory and homeostatic mechanisms could promote diversity among plastic responses in gene expression without altering higher-level phenotypic traits such as morphology and behavior (Badyaev, 2018; Fischer, Ghalambor, & Hoke, 2016a; Renn & Schumer, 2013). Finally, because fish in low-predation habitats experience relaxed selection on predator-induced plasticity in conjunction with low effective populations sizes, some of the evolution of transcriptional plasticity we report likely arose as a product of genetic drift, rather than selection for altered plastic responses (Lynch, 2007). Documenting the evolution of expression plasticity is an important first step, and additional studies are needed to understand the ubiquity and evolutionary sources of these patterns.

### Transcriptomic signatures of population divergence in two independent evolutionary lineages

We identified many genes differentially expressed between high- and low-predation populations in each river drainage. The absolute number of population DE transcripts was smaller in the Aripo drainage as compared to the Quare drainage, but the number of developmental and interaction differences was greater in the Aripo drainage. Our analyses provided no evidence that technical differences in expression magnitude, expression variance, or sample size influenced these differences between datasets in the number of genes reaching statistical significance. We therefore propose that differences in DE gene number between drainages are biological in origin, as high- and low-predation populations in the Quare drainage show greater genetic divergence than those in the Aripo drainage (Willing et al., 2010). Indeed, we note that the larger number of genes in the Quare dataset come from a large difference in the number of genes with population, but not rearing or interaction, effects. In sum, we suggest that the difference in the number of differentially expressed transcripts between datasets is primarily driven by the greater degree of genetic divergence between high- and low-predation populations within the Quare as compared to the Aripo river drainage (Willing et al., 2010).

When we compared DE gene sets between drainages, we found a small, but significant, number of population and rearing DE genes shared across lineages. However, the majority of DE genes were non-shared across drainages for both population (Aripo: 78%; Quare: 96%), and rearing (Aripo: 94%; Quare: 92%) effects, and we note that this pattern persisted at less stringent p-values cutoffs. Rearing DE genes showed a significant association in expression direction across lineages, with 67% of genes concordantly differentially expressed between lineages. In contrast, genes associated with genetic divergence between high- and low-predation populations in both drainages showed no signature of expression concordance across lineages. We suggest that these patterns across drainages point to a small number of core genes that exhibit predictable, plastic expression responses upon colonization of low-predation environments, but that lineage-specific selection pressures, differences in genetic background, non-adaptive processes (e.g. drift, inbreeding, founder effects), and alternative gene expression responses give rise to largely non-overlapping, non-concordant expression differences associated with parallel phenotypic adaptation.

A previous study in guppies performed a similar comparison of gene expression changes associated with adaptation to low-predation environments (Ghalambor et al., 2015) and found a strong signal of concordant differential expression in genes differentially expressed based on population of origin. Whereas the present study compared long-term natural population divergence across drainages, Ghalambor et al. (2015) characterized the early stages of adaptation of experimentally introduced low-predations populations derived from founders from a single high-predation source population within the same drainage. These contrasting findings in comparisons of population pairs within the same drainage versus across genetically distinct drainages highlight the impacts of standing genetic variation within the source population on mechanisms of divergence (Feiner, Rago, While, & Uller, 2017; Thompson, Osmond, & Schluter, 2019), particularly at early stages of evolution (Barrett & Schluter, 2008): while alternative transcriptional ‘solutions’ are possible, shared genetic background appears to bias evolutionary outcomes toward shared patterns. These findings are exciting as they extend to a mechanistic level the growing recognition of and interest in the non-parallel aspects of parallel adaptation, even in systems considered classical examples of parallel/convergent evolution (Bolnick et al., 2018; Manousaki et al., 2013; Paccard et al., 2020; Stuart et al., 2017).

Both adaptive and non-adaptive processes may contribute to the combination of shared and distinct transcriptional mechanisms we find associated with parallel, adaptive life-history, morphological, and behavioral phenotypes across lineages in guppies. First, as described above, differences in standing genetic variation likely influence which mechanisms are available to selection in response to common environmental conditions in different drainages (Barrett & Schluter, 2008; Thompson et al., 2019). Second, low-predation populations are typically established by a small number of individuals (Barson et al., 2009; Fraser et al., 2015; Willing et al., 2010), making them susceptible to the unpredictable, non-adaptive influences of founder effects, genetic drift, and/or inbreeding on gene expression divergence; although we note that work in three-spine stickleback suggests deviations from parallel evolution tend to result from selection rather than non-adaptive processes (Kaeuffer et al., 2011). Third, the large number of significantly evolved genes that did not overlap between drainages may also represent adaptive responses to drainage- or site-specific environmental factors other than predation (Fitzpatrick, Torres-Dowdall, Reznick, Ghalambor, & Chris Funk, 2014; Zandonà et al., 2011). Finally, compensatory or homeostatic gene expression responses may arise in response to any of the above factors, leading to alternative transcriptional configurations associated with similar higher-level phenotypes.

In sum, we suggest that genetic similarity among ancestral populations may channel low-predation populations within the same drainage toward shared transcriptional solutions (as in Ghalambor et al. 2015), while differences in standing genetic variation, drainage-specific environmental conditions, founder effects, and alternative compensatory changes can result in distinct mechanistic paths to arrive at shared organism-level phenotypes. We cannot definitely distinguish causal from non-adaptive and compensatory gene expression differences under these scenarios – indeed, it is likely a combination of these factors that contribute to distinct transcriptional patterns associated with parallel adaptation. Nonetheless, alternative transcriptional patterns suggest that mechanistic flexibility and ‘many-to-one’ mapping of gene expression to organism level phenotypes may facilitate adaptation.

### Conclusions

We assessed the extent to which adaptation to common environments targets predictable changes in transcript abundance across independent evolutionary events. Within lineages, genes with greater expression plasticity were more likely to diverge in expression between populations, but in the direction opposite of developmental changes. Despite a small number of differentially expressed genes shared across drainages, parallel adaptation to low-predation environments in independent lineages was associated with divergence in largely non-overlapping transcripts. While identification of shared genes is generally used as the starting point for work exploring mechanisms of parallel adaptation – and our work here indeed suggests that a small number of core genes may indeed be critical during the earliest stages of adaptation – we propose that parallel evolutionary transitions are not limited to a small set of possible transcriptional mechanisms in guppies. Instead, our results highlight the potential for alternative transcriptional solutions associated with parallel, adaptive trait evolution even within a single species. Transcriptional network versatility, in which diverse alterative network configurations can produce common network outputs and behavioral phenotypes, may allow underlying networks to simultaneously accommodate the influences of selection, drift, and genetic background and thereby facilitate evolution in a species known for rapid adaptation to novel environments.

## Acknowledgements

We thank the members of the Colorado State University Guppy Group for fish care, especially Sarah E Westrick and Kimberly E Dolphin for help with tissue collection. We thank Laura R Stein for assistance with tissue processing and Cameron K Ghalambor for fruitful discussions and comments on earlier versions of the manuscript. All high-powered computing was performed on the Odyssey computing cluster supported by the FAS Science Division Research Computing Group at Harvard University. This work was supported by the National Science Foundation DDIG-1311680 (to EKF), RCN IOS-1256839 (to EKF), IOS-1354755 (to KLH), IOS-1354775 (to KAH), IOS-0934451 (to KAH), DEB-0846175 (to CKG), IIS-1545994 (to WZ), and US Department of Energy DE-SC0018344 (to WZ).

## Author contributions

EKF and KLH conceived of the study; EKF collected samples and performed molecular work; EKF and KAH performed gene expression mapping, transcript abundance estimation, and preliminary differential expression analyses; YS and WZ devised and performed statistical analyses with input from EKF and KLH; EKF and KLH wrote the manuscript with contributions from all authors. All authors gave final approval for publication.

## Data Accessibility

Raw sequencing reads are available through the NCBI SRA repository (PRJNA601479). R code for statistical analyses is available on GitHub.

## Glossary

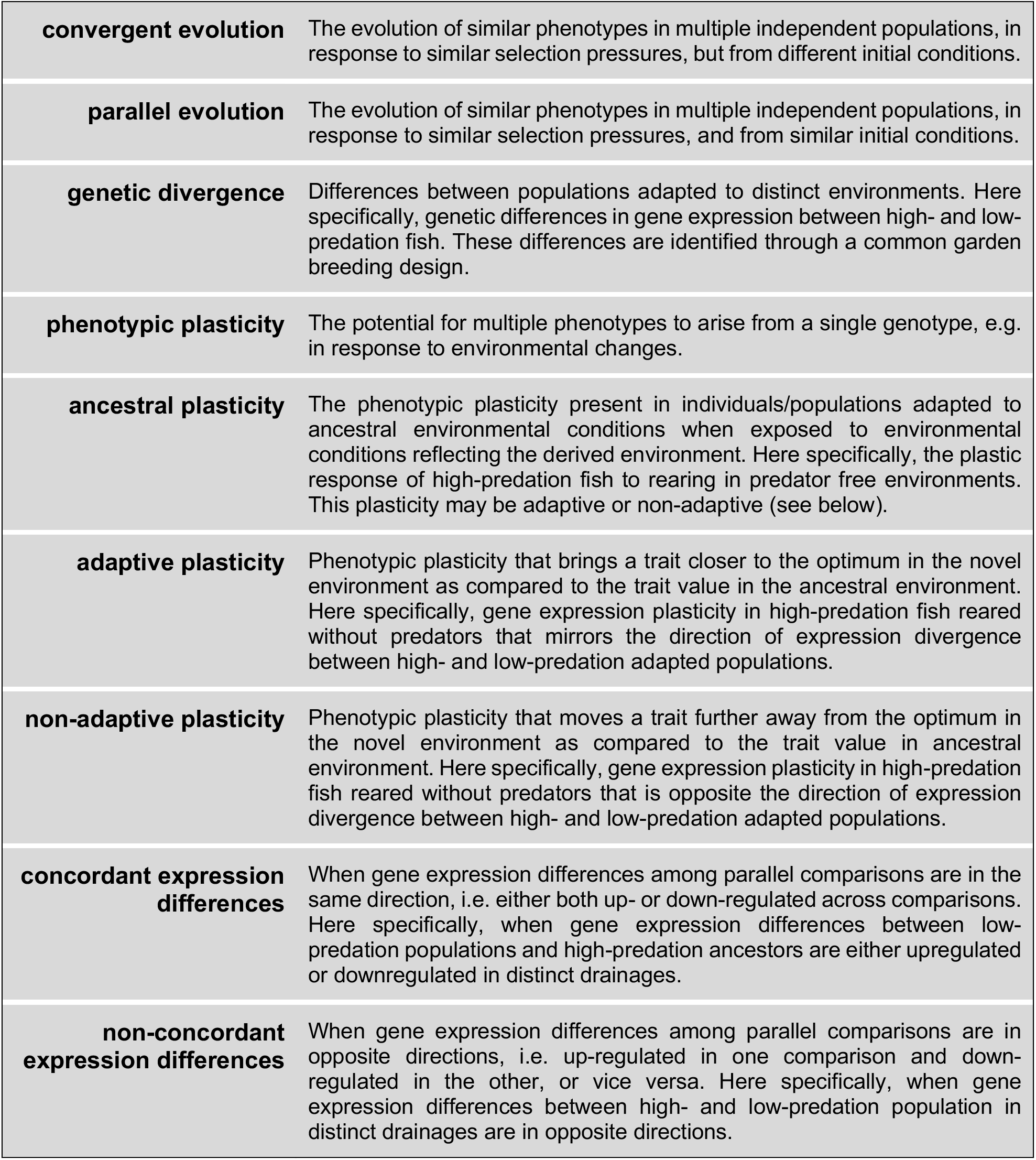

## Supplemental Information

**Table S1.** Transcripts filtered from the analysis following identification as retinal contaminants. Provided as Excel file.

**Table S2**. Transcripts significantly differentially expressed between high- and low-predation populations in the Aripo drainage. Provided as Excel file.

**Table S3**. Transcripts significantly differentially expressed between high- and low-predation populations in the Quare drainage. Provided as Excel file.

**Table S4**. Results of GO enrichment analyses for population, rearing, and interaction effects in the Aripo drainage. Provided as Excel file.

**Table S5**. Results of GO enrichment analyses for population, rearing, and interaction effects in the Quare drainage. Provided as Excel file.

**Table S6**. Population DE transcripts overlapping between Aripo and Quare drainages. 185 transcripts were DE in both drainages, of these 89 had expression differences in the same direction in both datasets (concordant differential expression). Provided as Excel file.

**Table S7**. Rearing DE transcripts overlapping between Aripo and Quare drainages. 16 transcripts were DE in both drainages, of these 7 had expression differences in the same direction in both drainages (i.e. concordant differential expression). Provided as Excel file.

**Table S8**. Pst estimates for the Aripo and Quare drainages.

---

**Figure S1.**
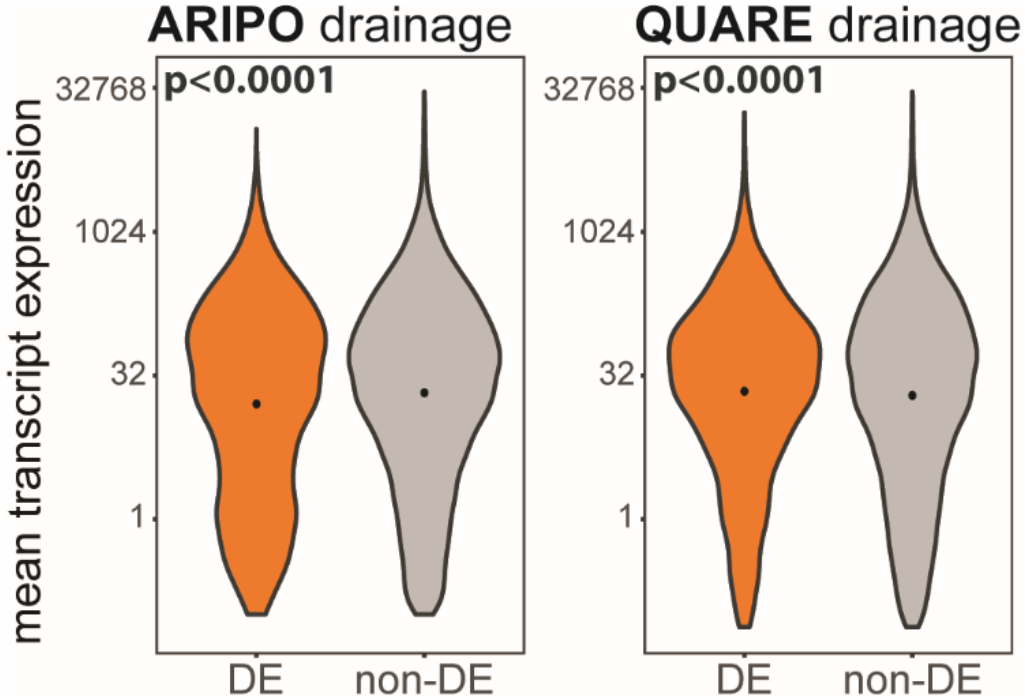
Differences in means but not variance of differentially expressed (DE) and non-differentially expressed (non-DE) transcripts. DE transcripts had significantly higher means in the Aripo drainage and lower means in the Quare drainage as compared to non-DE transcripts. Due to the large range of expression values, mean differences are plotted on a log scale (y-axis).

**Figure S2.**
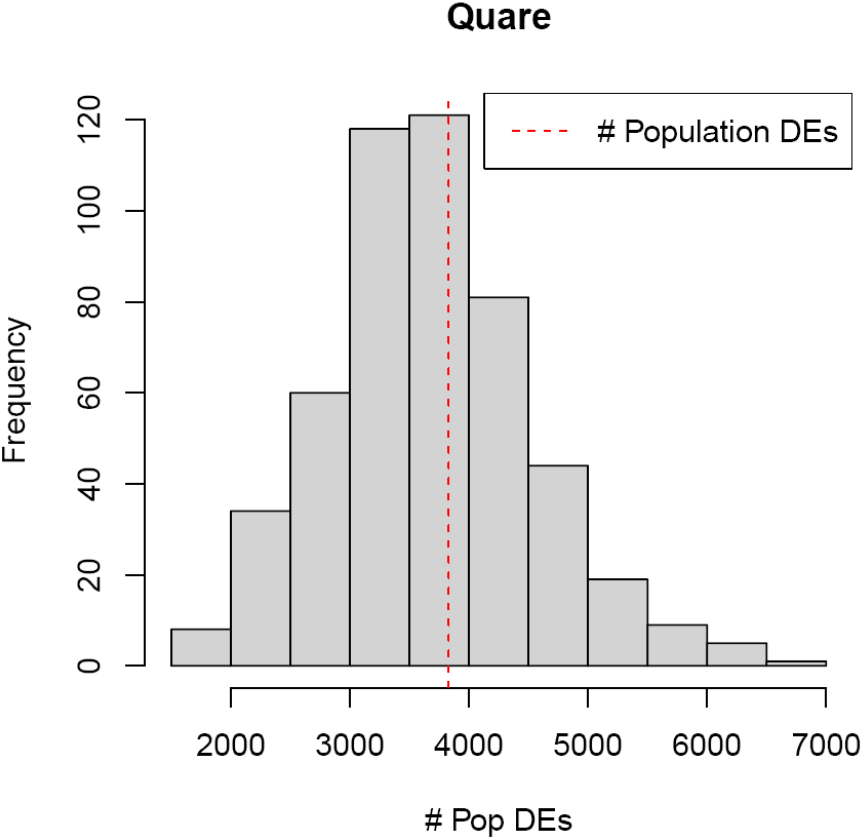
Outcome of down-sampling simulations. The Quare dataset was randomly down-sampled to generate a sample size equivalent to that of the Aripo dataset (40 samples, 10 per group). The histogram shows the distribution of the number of population DE genes obtained from 500 random down-samples. The number of DE genes in the full dataset (red dashed line) is not extreme compared to the distribution of the down-sampled datasets, indicating that sample size is not a critical factor for the difference in DE gene number between the two datasets.

