## Supplemental Methods for "Non-parallel transcriptional divergence during parallel adaptation"

This supplementary file details the statistical analysis used in the main paper. The pipeline (Figure 1) consisted of three steps: (1) pre-processing and normalization, (2) screening genes potentially dominated by retinal contamination of brain tissue, and (3) differential expression (DE) analysis. We analyzed RNAseq reads of brain tissue from guppies that came originally from two river drainages, Aripo and Quare rivers. We collected fish in each drainage from two populations, one low-predation and onw high-predation population (LP and HP, respectively). After two generations in the lab, we split siblings into two rearing conditions, either with (pred+) or without (pred-) predator chemical cues. We measured expression abundance of 23,254 genes in 40 brains from Aripo and 60 samples from Quare drainages. We ran separate models in the two river drainages due to differences in times of collection and methods of sample processing.

#### 1 Data filtering and screening

For data pre-processing, we filtered genes with low expression levels using the gene-wise overall counts as a criterion. For the Aripo drainage, we omitted 2685 genes with fewer than 50 counts total in the 40 samples, and in the Quare drainage, we filtered 1826 genes with fewer than 75 counts in the 60 samples. We performed principal component analysis (PCA) to identify samples that did not cluster with the bulk of the data. We excluded two samples from Quare, leaving 58 individuals total in the analysis.

Preliminary clustering analysis provided evidence for substantial retinal contamination in three samples in the Aripo dataset that could distort the subsequent DE analysis. We employed the function `estimateSizeFactors-ForMatrix()` in the `DESeq2` package in `R` to perform normalization separately within each drainage to account for variation in library sizes among samples. We then applied a screening procedure to detect potentially contaminated transcripts by inspecting the gene-wise total correlations to seven seed genes known to be highly expressed in the retina (rhodopsin, ENSPREG00000016208; red-sensitive opsins ENSPREG00000022016, ENSPREG00000022153 and ENSPREG00000008824; green-sensitive opsins ENSPREG00000002751 and ENSPREG00000002827; blue-sensitive opsin ENSPREG00000022188). Levels of all seven of these seed genes were high correlated in Aripo samples, so we expected other transcripts highly expressed in retinal tissues would also be positively correlated to the seed genes. We thus used large overall correlations to seed genes as evidence that expression levels of other genes were dominated by the retinal tissue contamination. We conducted this procedure only in the Aripo drainage, as the Quare drainage had uniformly low retinal contamination and hence provided little variation for detecting transcripts dominated by retinal tissue expression. For the $g$th gene, we tested $H_{0g}$ : the total correlation of the $g$th gene to seed genes is less than or equal to zero in the Aripo drainage. To do so, we applied the following procedure for testing the positive correlation using the sum of multiple Kendall's tau's [6] while controlling false discover rate (FDR) using the procedure from [2]. Given $G$ transcripts, $K$ seed genes, and $M$ groups (defined by the combination of population and rearing condition), the procedure is detailed as following.

1. Calculate the test statistic $\widehat{S}_g = (KM)^{-1/2} \sum_{k=1}^{K} \sum_{m=1}^{M} \widehat{\tau}_{gk,m}$ for $g = 1, \ldots, G$, $k = 1, \ldots, K$, and $m = 1, \ldots, M$, where $\widehat{\tau}_{gk,m}$ is the Kendall's tau between the $g$th gene and the $k$th seed gene in the $m$th group. This test statistic represents the average Kendall's tau for the focal gene and each seed gene calculated separately in all four groups.

2. Conduct the FDR controlling procedure based on the nonparametric bootstrap [3].

    (a) Sample with replacement within each group.

(b) Obtain bootstrap version of $\widehat{S}_g$,

$$\widehat{S}_g^{*,b} = \frac{1}{(KM)^{1/2}} \sum_{k=1}^{K} \sum_{m=1}^{M} \widehat{\tau}_{gk,m}^{*,b}$$

for $b = 1, \cdots, B$ bootstrap samples, where $\widehat{\tau}_{gk,m}^{*,b}$ is the Kendall's tau for the bootstrapped sample between the $g$th gene and the $k$th seed gene in the $m$th group .

(c) Set the mean proportion of rejections at a given threshold $t$ as $\theta_t^* = (BG)^{-1} \sum_{b=1}^{B} \sum_{g=1}^{G} \mathbb{I}(\widehat{S}_g^{*,b} > t)$, and calculate the test statistic threshold needed to reach the prespecified FDR correction $\alpha$ as

$$\widehat{t}_\alpha = \inf \left\{ t > 0 : \frac{\theta_t^* G}{\max\{\sum_{g=1}^{G} I(\widehat{S}_g \geq t), 1\}} \leq \alpha \right\}$$

3. Reject $H_{0g}$ and conclude the $j$th gene should be screened as likely retinal contaminant whenever $\widehat{S}_g > \widehat{t}_\alpha$.

Based on $2,000$ bootstrap iterations, we found $1,558$ potentially contaminated transcripts at the nominal FDR level $\alpha = 0.2$ to remove most spurious effects from the contaminants. We removed the potential retinal contaminants from both the Aripo and Quare drainages. The remaining analyses focused on the remaining $19,004$ and $19,902$ transcripts in the Aripo and Quare data, respectively.

### 2 Differential expression analysis

For the differential expression analysis, we fitted data to the generalized linear mixed model using `glmer.nb()` in `lme4` (R version 3.5.2, [1]). We considered a 2-way ANOVA model with interactions, which accounts for the partially crossed random effects due to week and family. The model is described as following. Let $Y_{ig,jl}$ be a negative binomial random variable with

$$\mathbb{E}\{Y_{ig,jl}\} = \exp\left\{\mu_g + \alpha_{g,j} + \beta_{g,l} + (\alpha\beta)_{g,jl} + (w)_{i_w} + (f)_{i_f}\right\}, \tag{1}$$

where $Y_{ig,jl}$ denotes the expression level of the $g$th gene from the $i$th sample for $i = 1, \ldots, n$, $j$ denotes the population (HP or LP), and $l$ denotes the rearing condition (pred+ or pred-). For the $g$th gene, $\alpha_{g,j}$, $\beta_{g,l}$, and $(\alpha\beta)_{g,jl}$ denote the main effect of population, the main effect of rearing,and their interactions, respectively; $w$ and $f$ are random effects due to week and family, respectively, while $i_w$ and $i_f$ are functions linking the family and week indices to the sample index. Implementation was carried by optimizers in `optimx` R package [8, 9] to avoid numerical instabilities.

We performed the differential expression analysis for the two main effects, four simple effects, and interaction. Corresponding $p$-values were obtained by the Wald test [5] and $q$-values [10] controlling the false discovery rate at 0.05 were obtained by `fdrtool` R package [11].

### 3 Inference on the relationship between plasticity and evolution

To understand whether ancestral plasticity influenced expression divergence, we compared conditional probabilities. The two simple effects of interest are the simple effect representing phenotypic plasticity in the ancestral population, which is estimated by $\widehat{\delta}_{1g} := -(\widehat{\beta}_{g,l} + \widehat{\alpha\beta}_{g,jl})$ based on model (1), and the population differences measured in the novel environment lacking predator cues, which is similarly estimated by $\widehat{\delta}_{2g} := \widehat{\alpha}_{g,j} + \widehat{\alpha\beta}_{g,jl}$ for the $g$th gene.

First, we asked whether a significant simple effect of rearing condition in the ancestral population (rearing effect in the HP population) promoted a significant simple effect of population in the novel environment (HP-LP population in the pred-environment). For that, we denote the probability of a significant population simple effect conditional on the significant and non-significant rearing effects, which are $\mathbb{P}$(population differences|phenotypic plasticity) and $\mathbb{P}$(population differences|no phenotypic plasticity) by $p_1$ and $p_2$, respectively. The hypothesis of interest is $H_0^{\mathrm{I}} : p_1 = p_2$. With estimates from model (1), we can estimate

$$\widehat{p}_1 = \frac{\sum_{g=1}^{G} \mathbb{I}\{\widehat{\delta}_{1g} \text{ and } \widehat{\delta}_{2g} \text{ are significant}\}}{\sum_{j=1}^{G} \mathbb{I}\{\widehat{\delta}_{1g} \text{ is significant}\}},$$

$$\widehat{p}_2 = \frac{\sum_{g=1}^{G} \mathbb{I}\{\widehat{\delta}_{2g} \text{ is significant, but } \widehat{\delta}_{1g} \text{ is not significant }\}}{\sum_{j=1}^{G} \mathbb{I}\{\widehat{\delta}_{1g} \text{ is not significant}\}},$$

and draw inferences about $H_0^{\mathrm{I}}$ using a bootstrap procedure detailed below.

We further analyzed an association between the directions of plasticity and evolution using two additional comparisons of conditional probabilities based on these simple effects. If the divergence measured in the novel pred-environment is in the same direction as the ancestral plasticity, then the two simple effects share common signs and $\delta_{1g}\delta_{2g} > 0$. On the other hand, if the expression divergence is opposite the direction of plasticity and hence partially or fully reverses the plasticity, then $\delta_{1g}\delta_{2g} < 0$. We asked whether divergence was more likely in the same or opposite directions 1) for all genes with a significant simple effect of population in the novel pred-environment and 2) for the subset of genes with significant simple effects of population that also had significant effects of rearing condition. Statistically, these two problems can be studied by comparing conditional probabilities. Problem 1 requires testing $H_0^{\mathrm{II}} : p_{\text{same},1} = p_{\text{opposite},1}$ using estimates

$$\widehat{p}_{\text{same},1} = \frac{\sum_{g=1}^{G} \mathbb{I}\{\widehat{\delta}_{1g}\widehat{\delta}_{2g} > 0, \widehat{\delta}_{2g} \text{ is significant}\}}{\sum_{g=1}^{G} \mathbb{I}\{\widehat{\delta}_{2g} \text{ is significant}\}},$$
$$\widehat{p}_{\text{opposite},1} = \frac{\sum_{g=1}^{G} \mathbb{I}\{\widehat{\delta}_{1g}\widehat{\delta}_{2g} < 0, \widehat{\delta}_{2g} \text{ is significant}\}}{\sum_{g=1}^{G} \mathbb{I}\{\widehat{\delta}_{2g} \text{ is significant}\}}.$$

Similarly, problem 2 requires testing $H_0^{\mathrm{III}} : p_{\text{same},2} = p_{\text{opposite},2}$ using estimates

$$\widehat{p}_{\text{same},2} = \frac{\sum_{g=1}^{G} \mathbb{I}\{\widehat{\delta}_{1g}\widehat{\delta}_{2g} > 0, \widehat{\delta}_{1g} \text{ and } \widehat{\delta}_{2g} \text{ are significant}\}}{\sum_{g=1}^{G} \mathbb{I}\{\widehat{\delta}_{1g} \text{ and } \widehat{\delta}_{2g} \text{ are significant}\}},$$
$$\widehat{p}_{\text{opposite},2} = \frac{\sum_{g=1}^{G} \mathbb{I}\{\widehat{\delta}_{1g}\widehat{\delta}_{2g} < 0, \widehat{\delta}_{1g} \text{ and } \widehat{\delta}_{2g} \text{ are significant}\}}{\sum_{g=1}^{G} \mathbb{I}\{\widehat{\delta}_{1g} \text{ and } \widehat{\delta}_{2g} \text{ are significant}\}}.$$

A naive approach to testing these hypotheses would be to plug in the estimates from model (1). That approach has two major shortcomings. First, it does not quantify the uncertainty of the estimates and therefore provides no statistical inference. Second, the plug-in estimators of these probabilities might be biased [4]. Though the bias correction is achievable for some simple hypotheses based on some canonical models, it is intractable for our models. The parametric bootstrap [3] provides a straightforward and valid way to draw inferences while correcting for potential biases [4]. Hence, we employed a parametric bootstrap-based procedure to draw inference on $H_0^{\mathrm{I}}$, $H_0^{\mathrm{II}}$, and $H_0^{\mathrm{III}}$:

1. Obtain the estimates of regression coefficients and the variance components of random effects based on the generalized linear mixed models (in the differential expression analysis).

2. Generate $B = 500$ bootstrap samples using the estimates from Step 1 and model (1) matching the sample numbers per group in the original dataset. For each $b = 1, \ldots, B$ bootstrap samples, re-do the differential expression analysis and compute the bootstrap statistics, $\widehat{p}_1^{(b)} - \widehat{p}_2^{(b)}$, $\widehat{p}_{\text{same},1}^{(b)} - \widehat{p}_{\text{opposite},1}^{(b)}$, and $\widehat{p}_{\text{same},2}^{(b)} - \widehat{p}_{\text{opposite},2}^{(b)}$ for $b = 1, \ldots, B$.

3. Construct 95% bootstrap confidence intervals via percentiles for $p_1 - p_2$, $p_{\text{same},1} - p_{\text{opposite},1}$, and $p_{\text{same},2} - p_{\text{opposite},2}$. For the intervals not contain 0, we reject the corresponding null hypotheses.

### 4 Calculation of gene wise $P_{\mathsf{ST}}$ values

To calculate $P_{\mathsf{ST}}$ for each transcript [7], we estimated variance components for population, family within population, and the residual variance using `SAS Proc Mixed` (version 9.4), after applying a log transformation to the normalized expression values described in main text. The models also included the fixed effect of rearing treatment and a variance component accounting for temporal variation (week of assay). We then calculated $P_{\mathsf{ST}}$ as

$$\frac{\sigma^2_{\text{population}}}{\sigma^2_{\text{population}} + 2h^2\{\sigma^2_{\text{Family(Population)}} + \sigma^2_{\text{Residual}}\}},$$

where the $\sigma^2_{\text{Subsript}}$ indicates the relevant variance component. We assumed that $h^2 = 0.5$, which produces a conservative (smaller) estimate of $P_{\mathsf{ST}}$ on the assumption that heritability of gene expression is likely to be $< 0.5$.

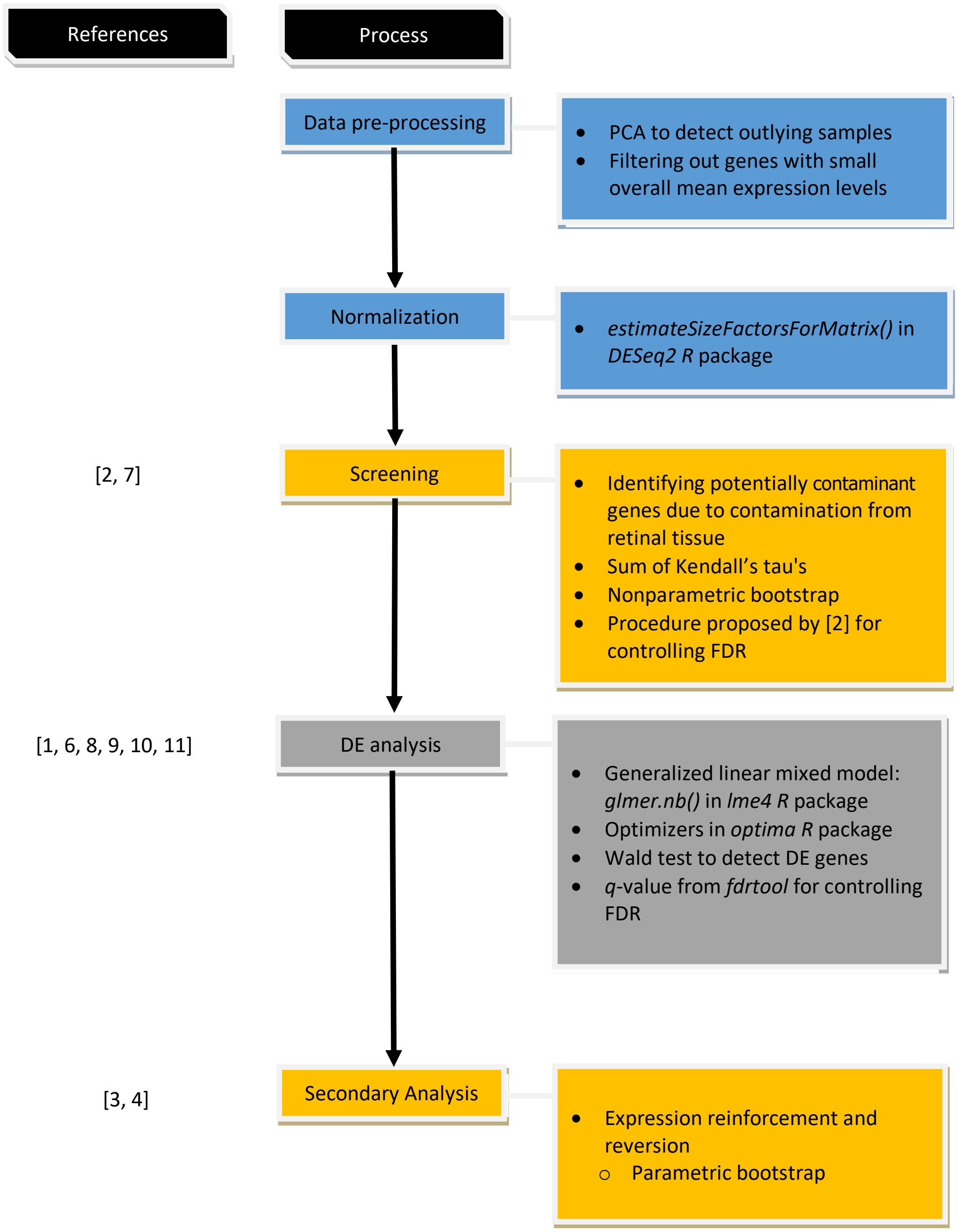


Figure 1: Workflow diagram for the statistical analysis.
